# Amalgams: data-driven amalgamation for the reference-free dimensionality reduction of zero-laden compositional data

**DOI:** 10.1101/2020.02.27.968677

**Authors:** Thomas P. Quinn, Ionas Erb

## Abstract

In the health sciences, many data sets produced by next-generation sequencing (NGS) only contain relative information because of biological and technical factors that limit the total number of nucleotides observed for a given sample. As mutually dependent elements, it is not possible to interpret any component in isolation, at least without normalization. The field of compositional data analysis (CoDA) has emerged with alternative methods for relative data based on log-ratio transforms. However, NGS data often contain many more features than samples, and thus require creative new ways to reduce the dimensionality of the data without sacrificing interpretability. The summation of parts, called amalgamation, is a practical way of reducing dimensionality, but can introduce a non-linear distortion to the data. We exploit this non-linearity to propose a powerful yet interpretable dimension reduction method. In this report, we present data-driven amalgamation as a new method and conceptual framework for reducing the dimensionality of compositional data. Unlike expert-driven amalgamation which requires prior domain knowledge, our data-driven amalgamation method uses a genetic algorithm to answer the question, “What is the best way to amalgamate the data to achieve the user-defined objective?”. We present a user-friendly R package, called amalgam, that can quickly find the optimal amalgamation to (a) preserve the distance between samples, or (b) classify samples as diseased or not. Our benchmark on 13 real data sets confirm that these amalgamations compete with the state-of-the-art unsupervised and supervised dimension reduction methods in terms of performance, but result in new variables that are much easier to understand: they are groups of features added together.

## 1 Introduction

Compositional data are a kind of relative data in which each part is only interpretable relative to the other parts [1, 5]. In the health sciences, many data sets produced by next-generation sequencing (NGS) have this property because of biological and technical factors that limit the total number of nucleotides observed for a given sample (often called the “constant-sum constraint”) [14, 15, 27, 20, 19, 38, 6]. As mutually dependent elements, it is not possible to interpret any component in isolation (at least without invoking the often untestable assumptions that underpin data normalization). The field of compositional data analysis (CoDA) offers an alternative way to analyze relative data by using log-ratio transforms. These transformations use one or more references to recast the data as log-contrasts [12]. The log-contrasts can then be analyzed using routine statistical methods, but must get interpreted as a ratio of the numerator parts to the reference denominator parts. Example log-ratio transformations include the additive log-ratio (alr) (which uses a single component as the reference) [1], the centered log-ratio (clr) (which uses the per-sample geometric mean as the reference) [1], and the isometric log-ratio (ilr) (which uses an orthonormal basis to define a set of arbitrary log-contrasts) [11].

Compositional data exist in a simplex with one fewer dimensions than parts. The ilr offers a theoretically ideal solution because its log-contrasts move the data from the simplex into real Euclidean space [11]. However, arbitrary log-contrasts lack interpretability. For example, how does an analyst make sense of the difference between the log of the product of two sets of parts, where each part is raised to a unique power? Balances were proposed as a more interpretable log-contrast, where each balance is a log-contrast between two geometric means [10]. An example 3-part balance is 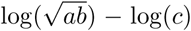. Indeed, Pawlowsky-Glahn et al. have shown how a set of “principal balances” can explain an ever-decreasing portion of the variance in analogy to principal components [34] (though principal balances can be correlated). Although more complicated than a simple log-ratio, having more parts means that a single balance can describe more variance than a single log-ratio. Balances have recently become popular for the analysis and classification of microbiome compositions [44, 46, 29, 40, 37].

Recently, Greenacre et al. have challenged the interpretability of balances [24]. We summarize the Greenacre et al. critique as follows: because the geometric mean depends on the ratios of the parts within, balances are not balances in the plain English sense of the word. Consider the balance between “*a* AND *b*” vs. “*c*” where *b > c*. We would expect that the “balance” would lean toward the combined weight of “*a*” and “*b*”. However, with a geometric mean, the balance will tip more toward “*c*” when “*a*” is rare. This is because the ilr balances are defined in log space. Instead of balances, Greenacre et al. proposed the summed log-ratio (SLR) as a more interpretable alternative [23]. An example 3-part SLR is log(*a* + *b*) − log(*c*). The summation of parts is called amalgamation, and Greenacre et al. encourage using domain-knowledge for amalgamation (i.e., *expert-driven amalgamation*) as a practical way of dealing with parts [22]. However, Egozcue & Pawlowsky-Glahn have criticized SLRs because, while scale-invariant, they are “non-linear functions in the Aitchison geometry of the simplex”, and so inter-sample distances can have “anomalous behavior” after amalgamation [12]. In summary, Greenacre et al. argue that summed log-ratios are an interpretable way to reduce the dimensionality of the data, while Egozcue & Pawlowsky-Glahn argue that summed log-ratios introduce a non-linear distortion to the data. Yet, non-linearity might be advantageous for situations in which the data need to be summarized in a non-trivial way. In this case, SLRs could provide a valuable addition to the compositional data analysis toolkit: an interpretable non-linear transform.

In this article, we propose *data-driven amalgamation* as a new method for reducing the dimensionality of compositional data. Unlike expert-driven amalgamation which uses domain knowledge, data-driven amalgamation uses an objective function. This objective function is user-defined for a given task, and combined with a search algorithm to answer the question, “What is the best way to amalgamate the data to achieve the objective?”. We show that data-driven amalgamation can be used to find a new 3-part simplex that efficiently visualizes the data according to any user-defined objective. We benchmark data-driven amalgamation across 13 health biomarker data sets, for two separate objectives: (a) to preserve a suitable distance between samples (where we consider three different measures), and (b) to classify samples as diseased or not. We show that the amalgamated features, which we call “amalgams”, can preserve inter-sample distances as well as principal components. Moreover, amalgams outperform principal components and principal balances as a feature reduction step before classification. We argue that amalgams are biologically meaningful concepts, and conclude the article by highlighting future areas of research.

## 2 Methods

### 2.1 Motivation

Greenacre et al. showed that using a pairwise log-ratio selection method in the presence of summed log-ratios does not necessarily distort inter-sample distances [23]. However, their example has two limitations. First, none of the “principal log-ratios” (i.e., the ones which explain the most variance) were summed log-ratios (SLRs). In other words, the SLRs happened to be the least important ratios. This raises the question, “What happens when the important log-ratios are summed log-ratios?”. Second, they only discuss expert-driven amalgamation. This raises another question, “Is it possible to replace expert-driven amalgamation with data-driven amalgamation?”.

While we do discuss SLRs in this article, we will focus on amalgamation more generally. Our motivation is to answer two research questions:

- Can we use a search heuristic to find an amalgamation that best preserves distance?
- Can we use a search heuristic to find an amalgamation that maximizes the prediction of a dependent variable?

The first question is an unsupervised machine learning problem that seeks to find a reduced feature space (i.e., a latent space) that accurately projects the data in fewer dimensions. The second question is a supervised machine learning problem. In both cases, amalgamation adds value over traditional dimension reduction methods because it makes the lower dimension features highly interpretable.

In this article, we benchmark data-driven amalgamation against the other dimension reduction methods used routinely for compositional data analysis, including principal components analysis (PCA), principal balance analysis (PBA) [34], and pairwise log-ratio selection (PRA) [21]. We define two tasks: (1) to obtain a compressed representation of inter-sample distances, and (2) to perform a feature reduction for binary classification. Figure 1 presents a schematic overview of our benchmark procedure.

**Figure 1:**
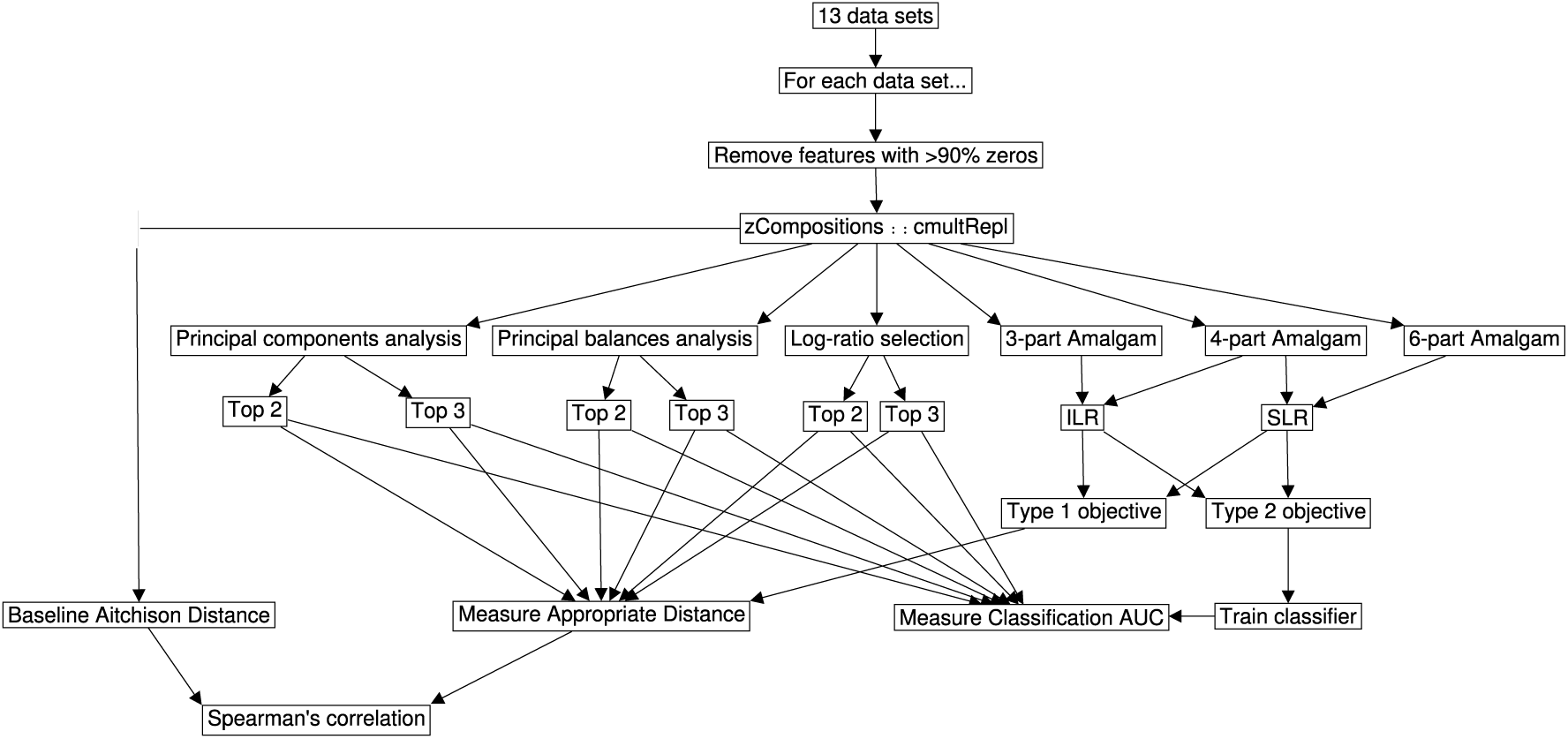
This figure presents an overview of the benchmark pipeline run for each data set. After feature removal and zero replacement, each data set underwent dimension reduction by principal components analysis (PCA), principal balance analysis (PBA), log-ratio selection (PRA), or data-driven amalgamation. The data-driven amalgams were then analyzed directly, or first converted to summed log-ratios. We used two criteria to benchmark the goodness of a dimension reduction method: (1) agreement between the baseline distances and the reduced-dimension distances and (2) accuracy of the reduced-dimension classifier.

### 2.2 Data-driven amalgamation

#### 2.2.1 The amalgamation matrix

An **amalgamation** is defined as the result of adding *D* components into *D*′ ≤ *D* mutually exclusive subsets [1]. A compositional data set **X** describing *N* compositions and *D* components can be amalgamated into a set of *D*′-part compositions via an *amalgamation matrix* **A**:

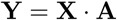

where *A*_*DD*′_ ∈ {0, 1} and 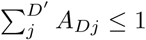.

The amalgamation has *D*′ new components which we call **amalgams**. The *amalgamation matrix* describes to which amalgam (as a column) the original component (as a row) belongs. Since one component should never contribute to more than one amalgam, the row sums of **A** is limited to 1 (when any 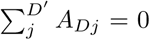, the amalgamation is also a sub-composition). Meanwhile, the column sums of **A** indicates how many components a single amalgam represents. Note that we use the term “amalgamation” to refer both to the amalgamation of the complete composition and to the amalgamation of a sub-composition. Figure 2 shows an example of amalgamation, and illustrates how one could conceptualize amalgamation as a feed-forward network.

**Figure 2:**
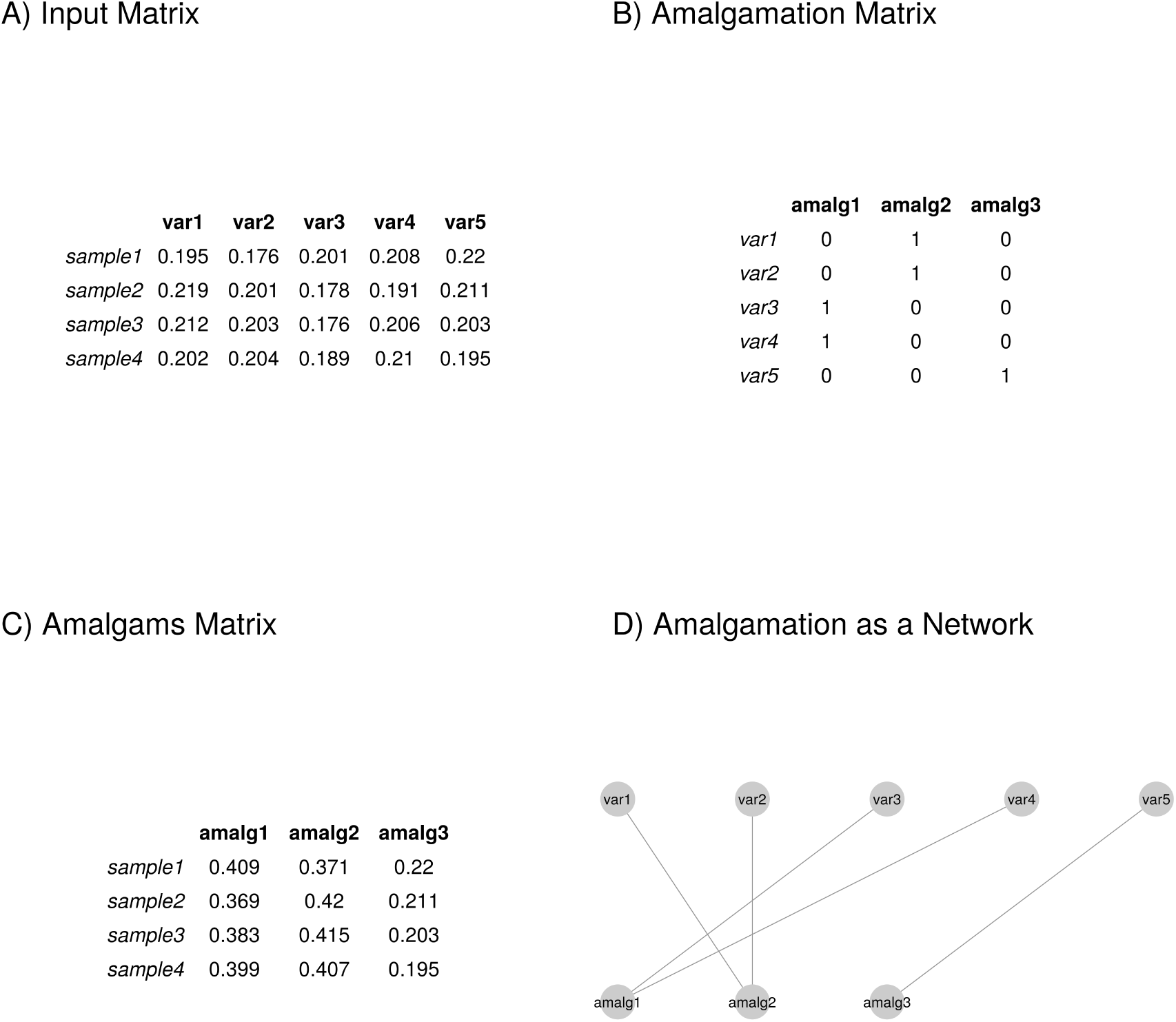
This figure shows an example amalgamation procedure, from the input compositions (Panel A) and the amalgamation matrix (Panel B) to the resultant amalgams (Panel C). The purpose of data-driven amalgamation is find the best *amalgamation matrix* for a given task. One could conceptualize amalgamation as a type of feed-forward network where each component has only one outgoing connection (Panel D). In this study, we search for an amalgamation matrix that maximizes an objective function; since this is equivalent to minimizing a loss, one could further conceptualize amalgams as a hidden layer in a (linearly activated) neural network. Although this suggests that we could use gradient descent, we choose to minimize the loss with a genetic algorithm because the amalgamation matrix is binary.

#### 2.2.2 The objective functions

Data-driven amalgamation seeks to find the best amalgamation matrix **A** for a given data set **X**:

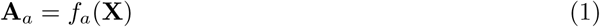

where *f*_*a*_ is chosen to optimize an arbitrary objective denoted by *a*. Here, we consider two kinds of objectives: unsupervised (Type 1) and supervised (Type 2) objectives.

#### 2.2.3 Type 1 objectives

Our Type 1 objectives seek to preserve the distance *d* between samples. This is an “unsupervised” objective that is designed for visualization tasks. We can express this objective in terms of maximizing the Pearson’s correlation *ρ* between the vectors of the original distances and the amalgamated distances:

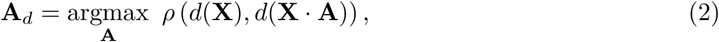

We consider 3 distances. First, we consider the log-ratio, or Aitchison, distance, which is the Euclidean distance obtained from clr-transformed data. It has a number of advantages, including scale invariance and sub-compositional dominance, that make it a preferred distance for compositional data. It can be defined as

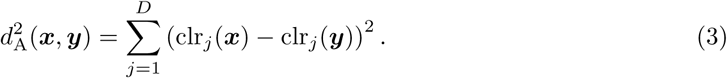

(Here we denoted the *j*-th component of the clr-transformed data by clr_*j*_.)

However, this may not be the most appropriate measure when considering amalgamations. Instead, we may want our distance measure to observe a natural continuity property: when we amalgamate parts that are identical in two samples, the distance between these samples should be unaffected by the merging of the parts. For compositional data, the notion of identity between parts across samples can be thought of loosely as a *proportionality* of the parts. Proportional parts have a vanishing log-ratio variance (i.e., they behave in an entirely coordinated way) [27]. Such parts are also known as “distributionally equivalent” [25].

Second, we consider the weighted Aitchison distance which, unlike its unweighted form, has distributional equivalence [25], meaning that it is unaffected by the merging of proportional parts. It can be defined as

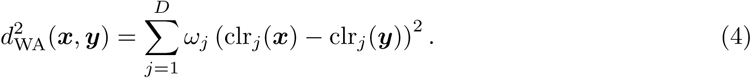

The exact form of the weights *ω*_*j*_ is not important, and we will use the simplest possibility by weighting each part by the total sum of all counts for that feature (i.e., the column sum). Note that, when merging parts, their weights will be added too.

Third, we consider a distance based on information-theoretic considerations. A composition is formally equivalent to a vector of discrete probabilities, and thus we could apply the Shannon index *H*. Advantageously, this measure is also scale-invariant when compositions are normalized to 1. Amalgamations over indices *j* ∈ 𝒜 can then be considered a *coarse-graining* [8] of the parts. It is well known [43] that Shannon entropy can be expressed as the sum of the entropy of the parts resulting after coarse-graining and of the coarse-grained parts themselves, where the latter are renormalized and weighted by their sum:

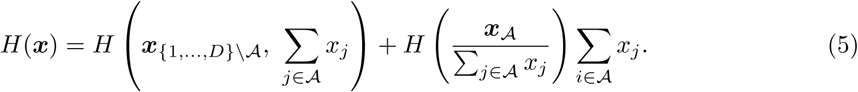

It is easy to show that the relative entropy

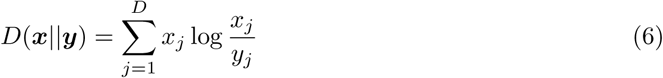

also remains invariant when merging distributionally equivalent parts (see Supplement). For simplicity, we consider the symmetrized version

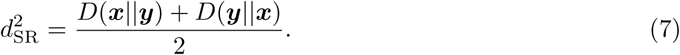

It is well known that the maximum-likelihood estimator of entropy is negatively biased for under-sampled data, e.g., [7]. To obtain a better empirical estimate of relative entropy from genomic data matrices, one could use the James-Stein type shrinkage estimator implemented in the R package `entropy` [26]. Since it allows for an estimate of the frequencies themselves, the shrinkage estimator can be used in conjunction with the other distance measures too. This approach is convenient because it naturally imputes the zeros that present a major problem for log-ratio analysis.

#### 2.2.4 Type 2 objectives

Our Type 2 objectives seek to maximize the percent of variance within the amalgamated data that is explained by a constraining matrix **L**. This is a “supervised” objective that is designed for prediction tasks. We can express this objective in terms of maximizing the relative size of the constrained eigenvalues of a (discriminant or) redundancy analysis (RDA) of ilr-transformed data:

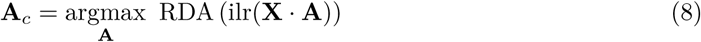

The ilr transformation implies an unweighted Aitchison distance for the RDA. Instead, any distance measure can be used via a square matrix of pairwise distances. These can be subjected to a classical multidimensional scaling to obtain principal coordinates that can be fed to the RDA.

#### 2.2.5 The summed log-ratio variant

We can create a set of summed log-ratios (SLRs) from an amalgamation by a function *f*_slr_ that takes the log of the first column of **Y** divided by the second column, the log of the third column divided by the fourth column, and so on:

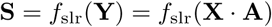

where the SLR matrix **S** contains *D*′*/*2 log-ratios.

Since SLRs have already moved the data out of the simplex, the Aitchison distance *d*_A_ (or its weighted version) is replaced with the Euclidean distance *d*_*E*_ in the Type 1 objectives:

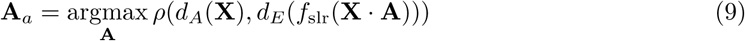

and the ilr-transformation is not performed in the Type 2 objectives:

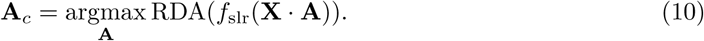

We do not use the summed log-ratios when evaluating relative entropies.

#### 2.2.6 The genetic algorithm

Since the *amalgamation matrix* is a binary matrix whose rows sum to 0 or 1, its parameters can be solved by a genetic algorithm with only *D* ∗ ceiling(log_2_(*D*′ + 1)) bits.

#### 2.2.7 Implementation

Here we present the `amalgam` package for the R programming language which solves the aforementioned objectives as a reproducible and easy-to-use software tool. Below, we show an example of its use for mock data.

~~~
*# install from GitHub*
devtools : : **install_**github (“tpq **/**amalgam”)
*# Load package and sample data*
**library** (amalgam)
**data** (iris)
*# find best amalgamation*
A **<**− amalgam (x = iris [, 1 : 4],
              n. amalgams = 3, *# how many amalgams to return*
              maxiter = 50, *# how long to run genetic algorithm*
              objective = objective. keep Dist, *# if preserving distance
              # objective = objective*. *keepWADIST # another distance
              # objective = objective*. *keepSKL # another distance
              # objective = objective*. *maxRDA, # if maximizing RDA*
              z = iris [, 5], *# only needed if maximizing RDA*
              asSLR = FALSE, *# if TRUE, n*. *amalgams must be even*
              shrink = FALSE) *# toggles James*−*Stein type shrinkage
# visualize results*
**plot** (A, **col** = iris [, 5])
~~~

The `x` argument defines the input data, the `n.amalgams` argument sets the number of amalgams, the `maxiter` argument sets the number of genetic algorithm iterations, the `objective` argument defines the objective, the `z` argument defines the constraining matrix, the `asSLR` argument toggles whether to convert the amalgams into summed log-ratios (SLRs), and the `shrink` argument toggles whether to use James-Stein type shrinkage. This package depends on the `GA` [42], `compositions` [45] and `vegan` [32] packages.

### 2.3 Benchmark evaluation

#### 2.3.1 Competing compositional methods

We benchmark amalgamation and summed log-ratios against competing dimension reduction methods designed for compositional data. This includes (1) a principal component analysis (PCA) of clr-transformed data [3] [implemented in `compositions` [45]], (2) a principal balance analysis (PBA) (using the log-ratio variance clustering heuristic) [34] [implemented in `balance` [36]], and (3) the pairwise log-ratio selection method proposed by [21] (PRA) [implemented in `propr` [39]]. For each dimension reduction technique, we consider the best 2 dimensions and the best 3 dimensions separately. Note that using 3 amalgams only occupies 2 dimensions because of the simplex.

#### 2.3.2 Data acquisition

We use the same 13 health biomarker data sets previously used to benchmark binary classification pipelines for compositional data [37]. These data sets were acquired from multiple sources [18, 31, 40, 41, 4, 9, 17, 47, 30] and span several difficult-to-study next-generation sequencing data types (including 16s, metagenomics, metabolomics, and microRNA). The number of samples, number of features, and outcomes-of-interest are described in Table 1. To facilitate between-study comparisons, these data underwent the same pre-processing steps as in [37]: for all data sets, we removed features that had more than 90% zeros; for the metabolomic and microRNA data sets, we only included features in the top decile of total abundance. The data are available already pre-processed for immediate use from https://zenodo.org/record/3378099.

**Table 1:** This table describes the data sets used to benchmark dimension reduction. Acronyms: CD Crohn’s disease; HC healthy control; MSM men who have sex with men; UC ulcerative colitis; IBD inflammatory bowel disease; CRC colorectal cancer. This table is reproduced from [37].

| Study Code | Source | Type | Features | Group 1 | Size | Group 2 | Size |
| --- | --- | --- | --- | --- | --- | --- | --- |
| 1a | selbal | 16s | 48 | CD | 662 | HC | 313 |
| 1b | selbal | 16s | 60 | MSM | 73 | Non-MSM | 55 |
| 2a | Franzosa et al. | Shotgun | 153 | IBD | 164 | HC | 56 |
| 2b | Franzosa et al. | Shotgun | 158 | CD | 88 | UC | 76 |
| 2c | Franzosa et al. | Metabolites | 885 | IBD | 164 | HC | 56 |
| 2d | Franzosa et al. | Metabolites | 885 | CD | 88 | UC | 76 |
| 3a | MicrobiomeHD | 16s | 278 | C. diff | 93 | Diarrhea | 89 |
| 3b | MicrobiomeHD | 16s | 610 | C. diff | 93 | HC | 154 |
| 3c | MicrobiomeHD | 16s | 1133 | CRC | 120 | HC | 172 |
| 3d | MicrobiomeHD | 16s | 1302 | CRC | 120 | Adenoma | 198 |
| 4a | TCGA | microRNA | 188 | Tumor | 1078 | Non-Tumor | 104 |
| 4b | TCGA | microRNA | 188 | Her2 | 77 | Non-Her2 | 927 |
| 4c | TCGA | microRNA | 188 | LumA | 524 | LumB | 194 |

#### 2.3.3 Zero replacement

All competing log-ratio methods fail in the presence of zeros. To address this impediment, we first replace zeros using the `cmultRepl` function from the `zCompositions` package [33]. Although we do not necessarily need zero replacement when using amalgamation or summed log-ratios, we use the zero-replaced data to make a fair comparison.

#### 2.3.4 Amalgams for preserving distance

The “gold standard” distance for compositions is the Aitchison distance [28, 2]. As such, we can evaluate the quality of a dimension reduction method based on how well the dimension-reduced distances agree with the true Aitchison distances [23]. For each data set, we measure this agreement as the Spearman’s correlation between the inter-sample distances from the dimension-reduced data and those from the full data. Note that when the reduced dimensions are already in log space, we compute a Euclidean distance instead.

We also compare how well each of our 3 dimension-reduced distances agrees with the corresponding baseline distances after 100 iterations, with and without James-Stein type shrinkage. This agreement is measured as Pearson’s correlation (as defined in the Type 1 objectives).

#### 2.3.5 Amalgams for classification

In a supervised setting, a good dimension reduction method should help classify a withheld test data. As such, we can evaluate the goodness of a dimension reduction method in terms of classification accuracy. For each data set, we measure classification accuracy by a cross-validation scheme in which we train a logistic regression classifier on a random sample of the dimension-reduced data. Then, the test set has its dimensions reduced *according to the training set rule*, thus ensuring test set independence. We repeat this procedure on 20 separate 67%-33% training-test set splits, and report the “out-of-the-box” performance without any hyper-parameter tuning because of the small sample sizes. The workflow is arranged using the `exprso` package [35].

## 3 Results and Discussion

### 3.1 Amalgams can preserve Aitchison distances

For NGS health biomarker data, each clinical sample is a composition. We can calculate inter-sample distances using the Aitchison distance. One critique against the use of amalgamation for dimension reduction is that it distorts inter-sample distances (i.e., it is not “sub-compositionally dominant”) [10, 12, 16]. Although amalgamation *can* distort distances, we perform data-driven amalgamation with an objective that preserves the Aitchison distances. Figure 3 shows the agreement between the baseline Aitchison distance and the distances computed using the top-2 or top-3 reduced dimensions. The x-axis presents 10 methods ranked from highest-to-lowest based on the (geometric) average agreement. Here, we see that the use of amalgamations (or summed log-ratios) preserves distances as well as a PCA, and both do better than a principal balance analysis of equal dimension. Supplemental Table 1 shows the 95% confidence interval for the median of the differences between each method, computed using the Wilcoxon Rank-Sum test.

**Figure 3:**
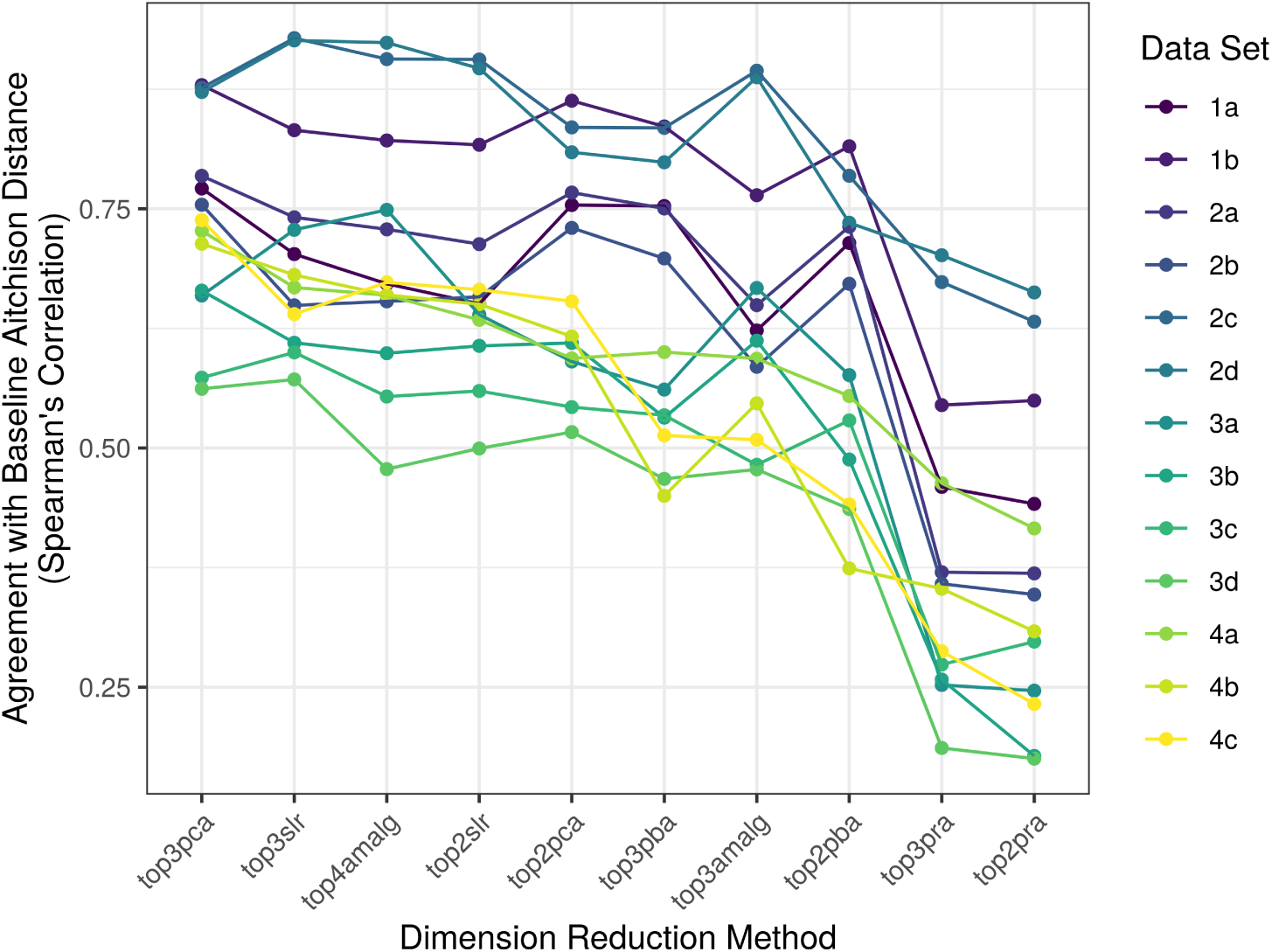
This figure shows the agreement between the baseline Aitchison distance and the distance computed on the dimension-reduced data (y-axis) for each method (x-axis), as grouped by the data set studied (color). Methods toward the left of the x-axis agree more with the baseline on average. Here, we see that using 4 amalgams or 3 summed log-ratios can actually preserve the inter-sample distances quite well. A statistical analysis of the differences is presented in Supplemental Table 1

Although data-driven amalgamation does not outperform PCA, it is arguably more interpretable. Although PCA is just a linear rotation of the data, its application to compositional data requires the use of clr-coordinates. Consequently, the coefficients of each principal component actually form a complex log-contrast where each variable (e.g., gene or microbe) is raised to an arbitrary power, then multiplied together. On the other hand, each amalgamation is a simple sum of parts, and therefore exists as a pooled construct that is intuitive to biologists (indeed, one might relate each amalgam to a “gene module” or a “bacteria community”). Advantageously, amalgamation allows the analyst to visualize the data in the same space that they exist: a simplex. As an example, Figure 4 shows the 3-part (2D) and 4-part (3D) amalgams computed for the Franzosa et al. microbiome data set. Based on Figure 3 and Supplemental Table 1, we know that the distances in these figures are as coherent as a PCA plot of equal dimension. Yet, the variables of the simplex (i.e., the corners of the triangle) are easily understood: they are groups of bacteria added together.

**Figure 4:**
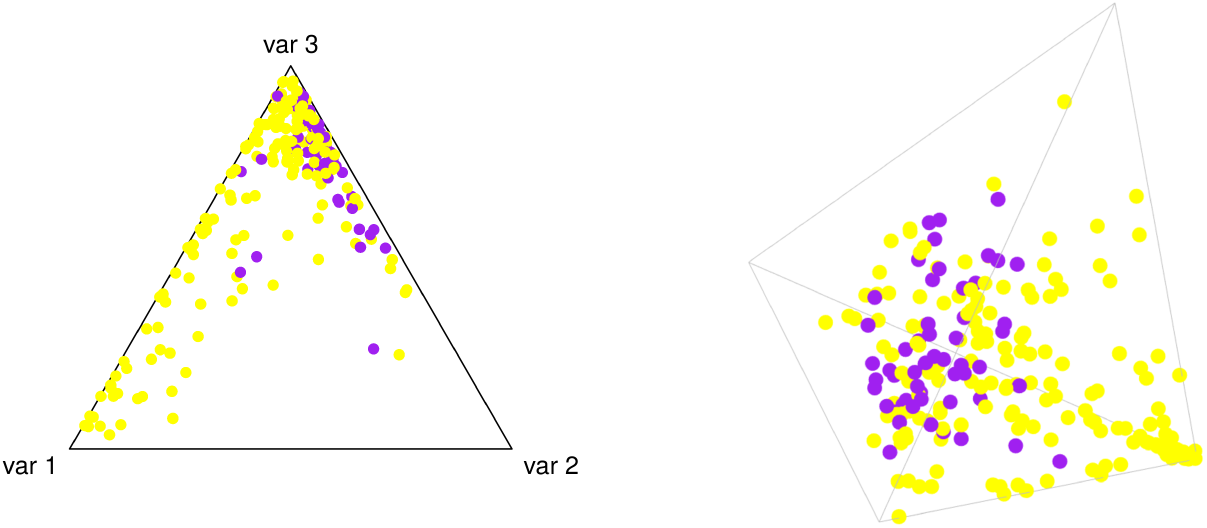
This figure projects the Franzosa et al. microbiome data across 3 amalgams (left panel) and 4 amalgams (right panel), chosen to preserve inter-sample distances. Light yellow dots show samples with inflammatory bowel disease, while dark purple dots show healthy samples. From Figure 3, we know that the distances in these figures are as coherent as a PCA plot of equal dimension. Yet, the variables of the simplex (i.e., the corners of the triangle) are easily understood, and allow the analyst to visualize the data in the same space that they exist: a simplex.

### 3.2 Amalgams can improve disease prediction

In the previous section, we show that data-driven amalgamation can successfully represent inter-sample distances in a coherent way (analogous to a principal coordinate analysis). By changing the objective function, we can instead search for an amalgamation that maximizes the separation between a binary class (analogous to a discriminant analysis). Supplemental Figures 1 and 2 show the area under the receiver operating curve (AUC) for classifiers trained on the top-2 or top-3 reduced dimensions (respectively), where each boxplot shows the distribution of AUCs across 20 randomly selected test sets. In both figures, we see the same trend: amalgams perform as well as or better than PCA, principal balances, and select log-ratios. Amalgams (and summed log-ratios) only under-perform on the Franzosa et al. data. Supplemental Table 2 shows the 95% confidence interval for the median of the differences between the AUCs for each method, computed using the Wilcoxon Rank-Sum test. Interestingly, the use of just 3 amalgams outperforms the use of 3 principal balances, the latter being a higher-dimensional representation.

As an example, Figure 5 shows the 4-panel output from the `amalgam` software. Notably, the bottom-right panel shows the distribution of samples across the 3-part simplex designed to maximize class separation. Visually, we can confirm that the amalgams do separate healthy guts from unhealthy guts based on the microbiome composition. However, amalgamation gives us a unique insight into the underlying process: the first amalgam not only associates with a sick gut, but makes up *most of the sick gut*. This perspective is reinforced by a boxplot of the per-sample pre-closure amalgam sums: the first amalgam takes up ∼10-60% of the entire gut composition of sick patients, compared with only ∼5% of healthy guts. Therefore, we can interpret the first amalgam as a kind of “sick gut community” whose members hardly ever appear in healthy patients.

**Figure 5:**
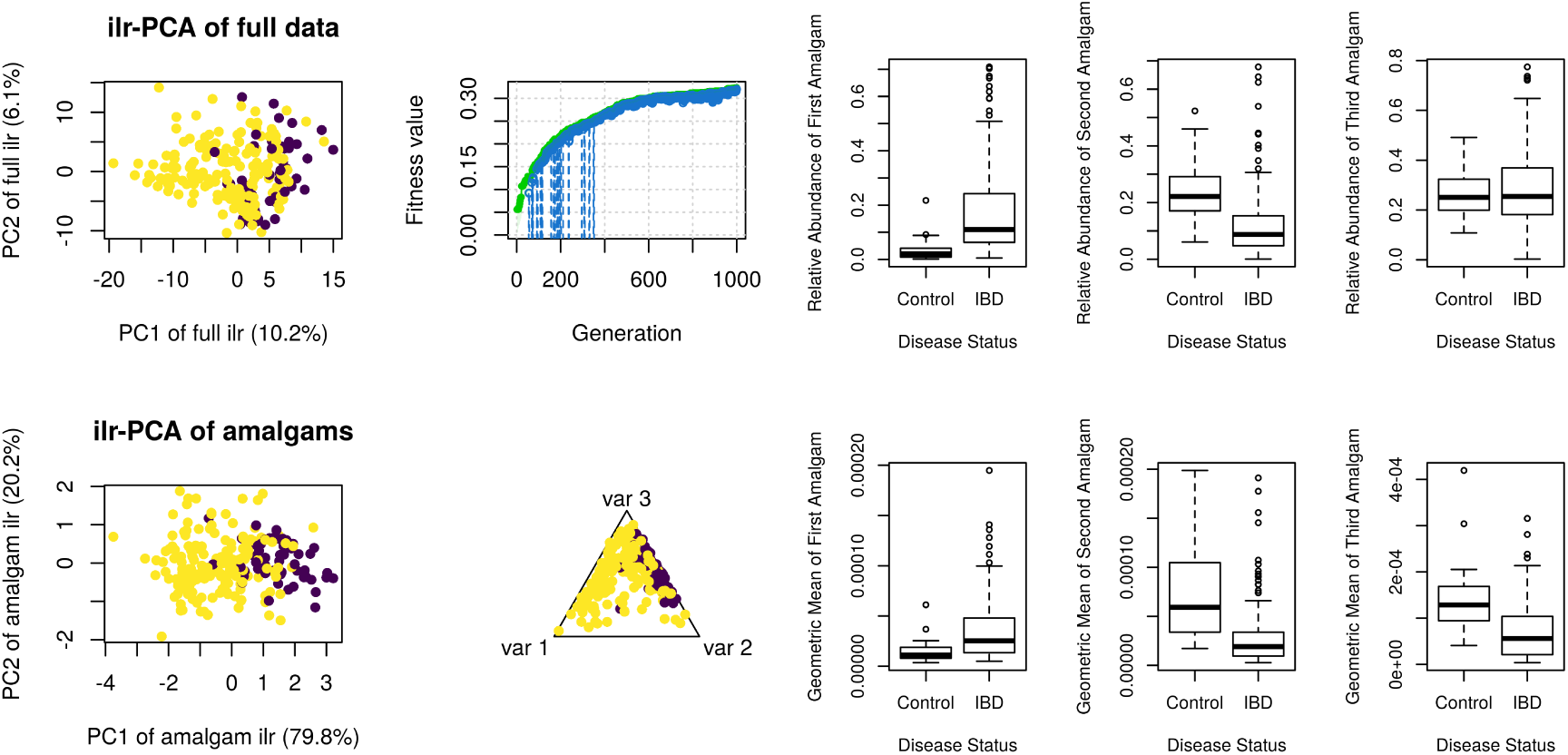
This figure shows the 4-panel output from the `amalgam` software, next to boxplots of the amalgam sums and their corresponding geometric means. The top-left panel shows a PCA of the ilr-coordinates. The top-right panel shows the “fitness” over each generation of the algorithm. The bottom-left panel shows a PCA of the ilr-coordinates of the amalgams. The bottom-right panel shows a ternary plot of all samples along the 3 amalgams selected to maximize the separation of sick guts (light yellow) and healthy guts (dark purple). The boxplots refer to these amalgams, where the per-sample sums are computed without a re-closure of the data (i.e., they are summed using the raw proportions). By only comparing the geometric means, one would miss the exciting insight that there exists a “sick gut community” signature that uniquely occupies up to one half of the sick gut.

Below the boxplots of the per-sample amalgams, we see boxplots of the per-sample geometric means. Just as amalgams are the building blocks of summed log-ratios, geometric means are the building blocks of balances. Although amalgamations and geometric means do not have to agree (as Greenacre et al. show in the beer-wine-spirit data), they do in this example. As such, misinterpreting a balance as if it were an amalgamation is of little consequence. However, by only comparing the geometric means, one would miss the exciting insight that there exists a “sick gut community” signature that uniquely occupies up to one half of the sick gut.

### 3.3 Amalgamation as an information bottleneck

The data benchmarked in this study were also benchmarked for other compositional data analysis classification procedures, including balance selection. Compared to these, we find that data-driven amalgamation performs as well as the balance selection method `selbal` [40], but under-performs when compared to using *all* ilr-transformed or clr-transformed features (see Table 2). This is not surprising when we consider that using only 3 (or 4) amalgams would have a limited capacity to explain the total structure of the data. In other words, it is possible that classifiers trained on so few amalgams *under-fit* the data. To test this hypothesis, we also trained logistic regression classifiers for *k* = [3, 6, …, 18, 21] amalgams. Figure 6 shows that 8 of the 13 data sets have a better classification AUC when using more amalgams. These findings reinforce the intuition that amalgams act as an “information bottleneck” when reducing the dimensionality of the data. If there are too few amalgams, the model will under-fit.

**Table 2:** This table reports the 95% confidence interval for the median of the difference between the amalgam-based logistic regression classifier AUCs and the other procedures benchmarked in [37]. Here, we see that amalgam-based classifiers perform as well as the balance selection method selbal [40], but under-performs when compared to using *all* ilr-transformed or clr-transformed features. Acronyms: SLR summed log-ratios; PBA principal balance analysis; ABA anti-principal balance analysis; RBA random balance analysis; DBA distal balance analysis; ACOMP raw proportions; CLR centered log-ratio transformed data.

|  | top3amalg vs. | top4amalg vs. | top2slr vs. | top3slr vs. |
| --- | --- | --- | --- | --- |
| Selbal | -0.0357 to 0.0033 | -0.0330 to 0.0048 | -0.031 to 0.007 | -0.023 to 0.014 |
| PBA | -0.055 to -0.014 | -0.053 to -0.013 | -0.051 to -0.011 | -0.0413 to -0.0031 |
| ABA | -0.053 to -0.012 | -0.052 to -0.010 | -0.0501 to -0.0096 | -0.0401 to -0.0014 |
| RBA | -0.057 to -0.016 | -0.055 to -0.014 | -0.053 to -0.013 | -0.043 to -0.005 |
| DBA | -0.07 to -0.03 | -0.069 to -0.027 | -0.065 to -0.027 | -0.056 to -0.017 |
| ACOMP | -0.018 to 0.024 | -0.018 to 0.026 | -0.016 to 0.027 | -0.007 to 0.035 |
| CLR | -0.064 to -0.023 | -0.062 to -0.021 | -0.059 to -0.020 | -0.049 to -0.012 |

**Figure 6:**
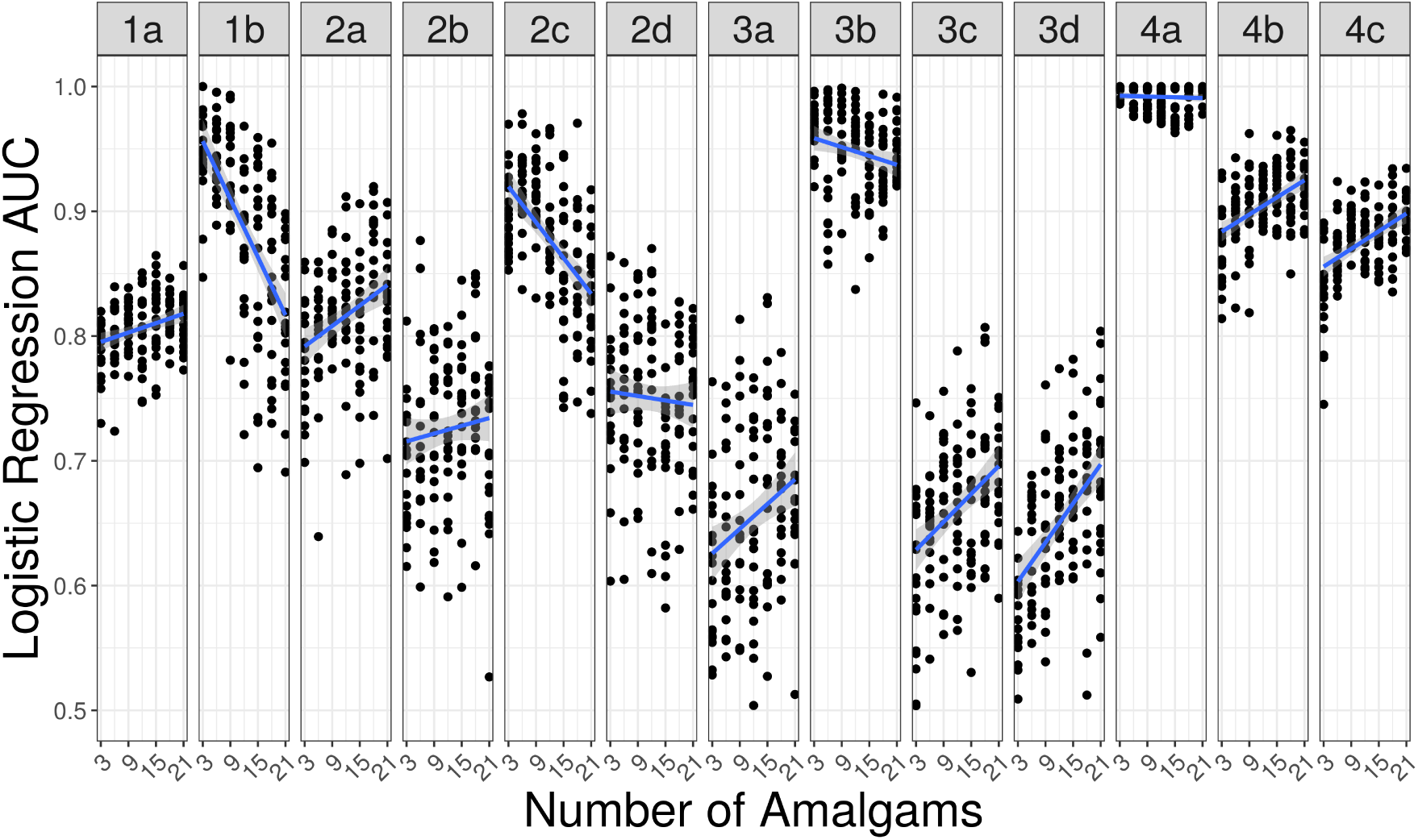
This figure shows the logistic regression classification AUC (y-axis) based on the number of amalgams (x-axis) used to train the model, organized by the data set under study (facet). Each point describes a different training-test set split, and a line is drawn to show the trend. Here, we see that 8 of the 13 data sets have a better classification AUC when using more amalgams. This might suggest that using 3 amalgams under-fit these data. Interestingly, 3 data sets do markedly worse with more amalgams. This might suggest that using more amalgams over-fit these data.

### 3.4 Alternatives to the Aitchison distance

Although the Aitchison distance has a number of advantages, including scale invariance and sub-compositional dominance, it may not be the most appropriate measure when considering amalgamations. Instead, we may want our distance measure to observe a natural continuity property: when we amalgamate parts that are identical in two samples, the distance between these samples should be unaffected by the merging of the parts. This property, called *distributional equivalence*, is found in the weighted version of the Aitchison distance [25], and also in relative entropy (see Supplement). Figure 7 shows how well each of these 3 dimension-reduced distances agrees with the corresponding baseline distances for 13 data sets. Here, we see that the distributionally equivalent distances tend to have better agreement after amalgamation. Better agreement means that the closest samples will remain close together while the furthest samples will remain far apart (though our use of Pearson’s correlation allows for any scale factor). This finding supports our hypothesis that distributionally equivalent distances are appropriate for amalgamated data. Interestingly, we see further improvement for all distances with James-Stein type shrinkage. This is noteworthy because for non-zero shrinkage, zeros are imputed directly, making it generally useful for zero-laden compositional count data like those encountered in microbiome or single-cell research.

**Figure 7:**
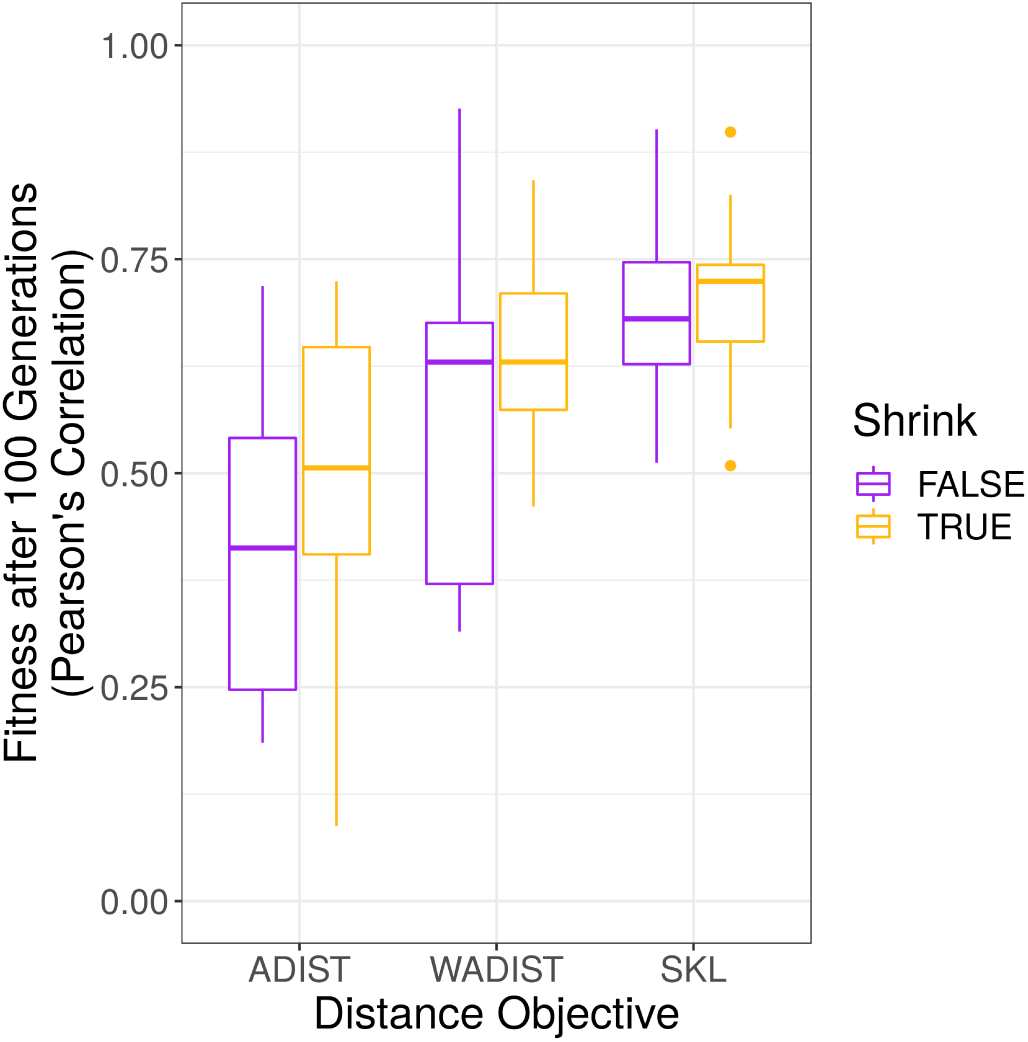
This figure shows how well the Aitchison (ADIST), weighted Aitchison (WADIST), and relative entropy (SKL) dimension-reduced distances agree with the corresponding baseline distances for 13 data sets (after 100 iterations, with and without James-Stein type shrinkage). Here, we see that the distributionally equivalent distances tend to have better agreement after amalgamation. We see further improvement with James-Stein type shrinkage.

## 4 Limitations and Future Directions

A major critique of amalgamation focuses on its non-linear behavior. However, we acknowledge this when designing our search heuristic, and instead use the non-linearity to our advantage. Still, data-driven amalgamation has some limitations. First, genetic algorithms, although faster than an exhaustive search, are still quite slow (especially when compared with a PCA). For example, the Franzosa et al. microbiome data starts to converge after ∼1500 iterations, which takes ∼3.5 minutes on an Intel i7 laptop computer. A PCA of an ilr of the same data set takes ∼50 milliseconds. Second, data-driven amalgamation appears less effective than other simple-but-fast methods for binary classification (e.g., the distal discriminative balance analysis method described in [37]), but easily generalized to multivariable regression. Third, data-driven amalgamation requires the user to select “hyper-parameters” to guide the dimension reduction, for example the number of amalgams. We see the importance of this hyper-parameter in Figure 6, where using too few (or too many) amalgams can impair classification accuracy. Fourth, amalgamation assumes that the relationship between the parts is explained by addition (a logical OR); as such, amalgamation would miss relationships explained by multiplication (a logical AND). On the other hand, balances would capture AND relationships but miss OR relationships.

It may be possible to resolve the first two limitations by relaxing the definition of the amalgamation matrix, for example by allowing the amalgamation matrix to become non-binary. This would allow an amalgam to equal the sum of parts of parts (not just the sum of parts). Although this expands the search space, it may enable the use of a faster search algorithm, such as gradient descent. Indeed, a total relaxation of all amalgamation matrix constraints would leave us with a single-layer neural network where the hidden layer is analogous to the amalgam set. We expect that re-factoring data-driven amalgamation as a neural network would improve the representative power of the amalgams and also increase the run-time. However, care is needed to define the weight constraints in a way that maintains the interpretability of the hidden layer. For example, the subtraction of parts would work arithmetically, but it is unclear what interpretation this would imply. One might maintain some interpretability by introducing explicit neural network constraints. For example, one might require that a component never contributes more than itself (i.e., 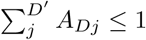), or that a component always contributes positively (i.e., 0 ≤ *A*_*DD′*_).

An important property that we have only mentioned in passing is the implicit handling of zeros that is achieved by amalgamation. We could envision objective functions that remove zeros across samples by specifically merging zero-laden parts with other parts. Another problem that can be alleviated by amalgamation is under-sampling: the merging of parts is a way of reducing dimensions without discarding data and can reduce the number of variables such that an inversion of their covariance matrix becomes possible. This enables, for example, an evaluation of partial correlations [13] on the new variables without having to resort to regularization.

## 5 Summary

In this report, we present data-driven amalgamation as a new method and conceptual framework for reducing the dimensionality of compositional data. Although amalgamation is criticized for distorting inter-sample distances, we show that data-driven amalgamation can preserve inter-sample distances as well as PCA when guided by an objective function. We also show that data-driven amalgamation can outperform both PCA and principal balances as a feature reduction method for classification, and performs as well as a supervised balance selection method called `selbal`.

Amalgamation not only allows the analyst to visualize the data in a lower-dimensional simplex (resembling the one in which the data naturally exist), but can also reveal interesting patterns about the relative abundances of the compositions. We demonstrate this through the discovery of a “sick gut community” bacterial signature that occupies more than one half of the sick gut, but is rarely found in healthy samples. We encourage principled research into data-driven amalgamation as a tool for understanding high-dimensional compositional data, especially zero-laden count data for which standard log-ratio transforms fail.

