## Supplement for "Amalgams: data-driven amalgamation for the reference-free dimensionality reduction of zero-laden compositional data"

### 1 Distributional Equivalence of Relative Entropy

Let us consider two compositions $\boldsymbol{x}$, $\boldsymbol{y}$ describing the same $D$ parts, which sum to 1 and are strictly greater than zero in both cases. In addition, let $x_{j^*} = mx_j$ and $y_{j^*} = my_j$, with $m$ some positive real number and $j$, $j^*$ indicating two parts of the compositions. By this definition, the parts are proportional (or *distributionally equivalent*) across the two samples . The relative entropy (or Kullback-Leibler divergence) between the compositions is then

$$\begin{aligned} D(\boldsymbol{x}||\boldsymbol{y}) &= \sum_{i=1}^{D} x_i \log \frac{x_i}{y_i} = \sum_{i\neq j,j^*} x_i \log \frac{x_i}{y_i} + x_j \log \frac{x_j}{y_j} + x_{j^*} \log \frac{x_{j^*}}{y_{j^*}} \\ &= \sum_{i\neq j,j^*} x_i \log \frac{x_i}{y_i} + x_j \log \frac{x_j}{y_j} + mx_j \log \frac{mx_j}{my_j} = \sum_{i\neq j,j^*} x_i \log \frac{x_i}{y_i} + (1+m)x_j \log \frac{x_j}{y_j} \\ &= \sum_{i\neq j,j^*} x_i \log \frac{x_i}{y_i} + (1+m)x_j \log \frac{(1+m)x_j}{(1+m)y_j} = \sum_{i\neq j,j^*} x_i \log \frac{x_i}{y_i} + (x_j + x_{j^*}) \log \frac{x_j + x_{j^*}}{y_j + y_{j^*}} \\ &= D((\boldsymbol{x}_{\{1,\ldots,D\}\setminus\{j,j^*\}}, x_j + x_{j^*})||(\boldsymbol{y}_{\{1,\ldots,D\}\setminus\{j,j^*\}}, y_j + y_{j^*})). \end{aligned} \quad (1)$$

Thus, relative entropy remains invariant after amalgamating the proportional parts. Clearly, the property of distributional equivalence carries over to a symmetrized version of relative entropy.

### 2 Supplemental Tables and Figures

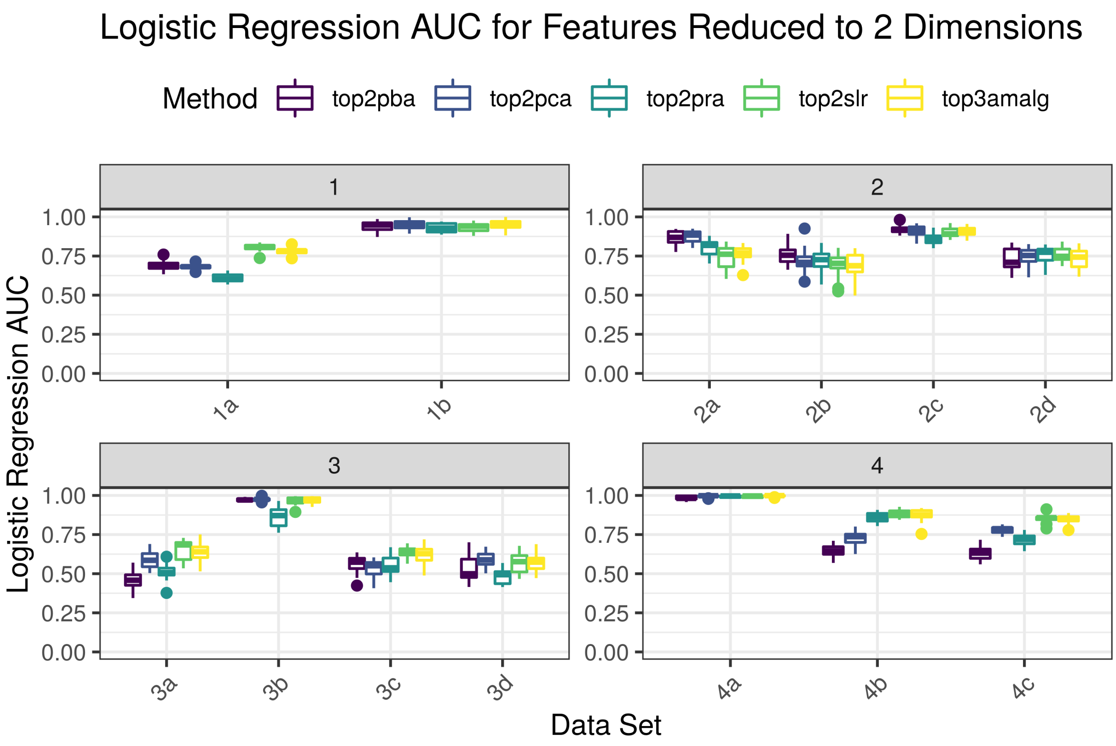


Figure 1: This figure shows the logistic regression classifier AUC (y-axis) for each data set (x-axis) separated by the dimension reduction method (color). Each method shown here reduces the data to 2 dimensions. Boxplots show the variability of performance over multiple training-test set splits. A statistical analysis of the differences between methods is presented in Table 2.

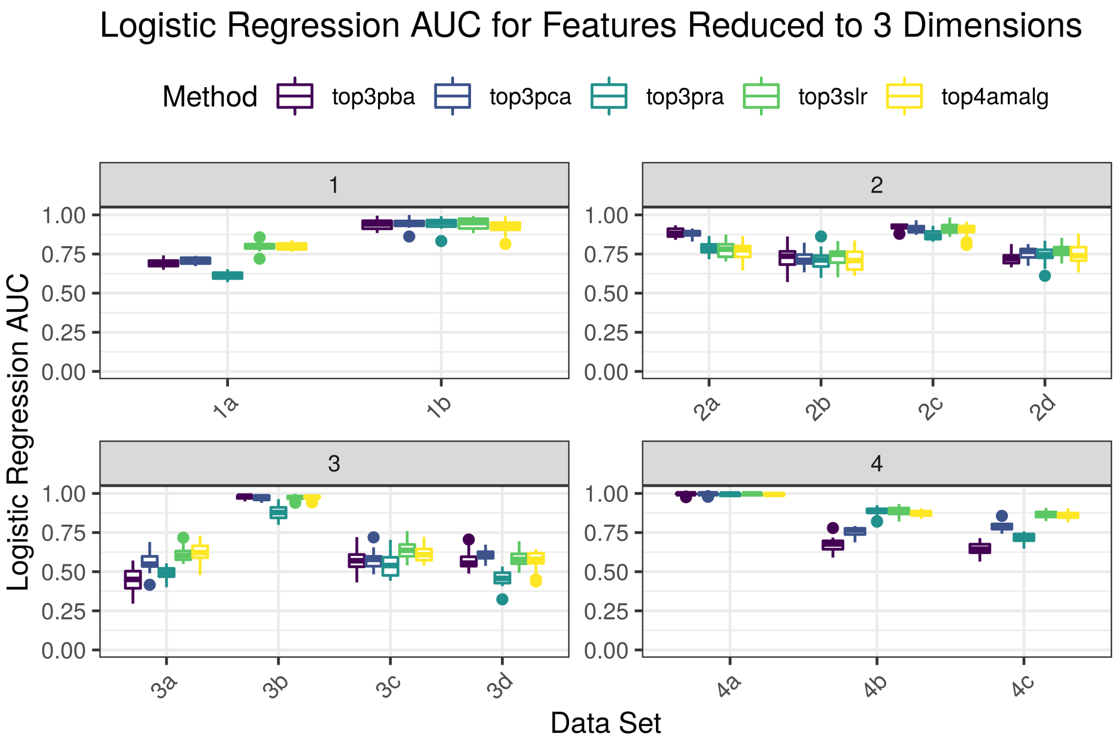


Figure 2: This figure shows the logistic regression classifier AUC (y-axis) for each data set (x-axis) separated by the dimension reduction method (color). Each method shown here reduces the data to 3 dimensions. Boxplots show the variability of performance over multiple training-test set splits. A statistical analysis of the differences between methods is presented in Table 2.

Table 1: This table compares how well each dimension reduction method preserves the baseline distances. Each entry shows the 95% confidence interval (CI) for the median of the differences between each method, computed using the Wilcoxon Rank-Sum test (where CI units measure the difference in Spearman's correlation). An all-positive or all-negative confidence interval suggests that the difference is significant at an unadjusted p-value of $p < 0.05$.

| | top2pca vs. | top3pca vs. | top2pra vs. | top3pra vs. | top2pba vs. | top3pba vs. | top3amalg vs. | top4amalg vs. | top2slr vs. | top3slr vs. |
|---|---|---|---|---|---|---|---|---|---|---|
| top2pca | — | -0.04 to 0.15 | -0.4 to -0.2 | -0.4 to -0.2 | -0.19 to 0.05 | -0.15 to 0.08 | -0.16 to 0.05 | -0.1 to 0.1 | -0.11 to 0.09 | -0.08 to 0.12 |
| top3pca | -0.15 to 0.04 | — | -0.5 to -0.2 | -0.5 to -0.2 | -0.250 to -0.008 | -0.21 to 0.03 | -0.218 to 0.008 | -0.13 to 0.06 | -0.15 to 0.04 | -0.13 to 0.06 |
| top2pra | 0.2 to 0.4 | 0.2 to 0.5 | — | -0.1 to 0.2 | 0.1 to 0.4 | 0.1 to 0.4 | 0.1 to 0.4 | 0.2 to 0.5 | 0.2 to 0.4 | 0.2 to 0.5 |
| top3pra | 0.2 to 0.4 | 0.2 to 0.5 | -0.2 to 0.1 | — | 0.08 to 0.36 | 0.1 to 0.4 | 0.1 to 0.4 | 0.2 to 0.4 | 0.2 to 0.4 | 0.2 to 0.4 |
| top2pba | -0.05 to 0.19 | 0.008 to 0.250 | -0.4 to -0.1 | -0.36 to -0.08 | — | -0.09 to 0.16 | -0.1 to 0.2 | -0.05 to 0.22 | -0.07 to 0.19 | -0.03 to 0.21 |
| top3pba | -0.08 to 0.15 | -0.03 to 0.21 | -0.4 to -0.1 | -0.4 to -0.1 | -0.16 to 0.09 | — | -0.1 to 0.1 | -0.08 to 0.17 | -0.1 to 0.1 | -0.07 to 0.18 |
| top3amalg | -0.05 to 0.16 | -0.008 to 0.218 | -0.4 to -0.1 | -0.4 to -0.1 | -0.2 to 0.1 | -0.1 to 0.1 | — | -0.03 to 0.17 | -0.03 to 0.15 | -0.01 to 0.17 |
| top4amalg | -0.1 to 0.1 | -0.06 to 0.13 | -0.5 to -0.2 | -0.4 to -0.2 | -0.22 to 0.05 | -0.17 to 0.08 | -0.17 to 0.03 | — | -0.11 to 0.08 | -0.08 to 0.10 |
| top2slr | -0.09 to 0.11 | -0.04 to 0.15 | -0.4 to -0.2 | -0.4 to -0.2 | -0.19 to 0.07 | -0.1 to 0.1 | -0.15 to 0.03 | -0.08 to 0.11 | — | -0.06 to 0.11 |
| top3slr | -0.12 to 0.08 | -0.06 to 0.13 | -0.5 to -0.2 | -0.4 to -0.2 | -0.21 to 0.03 | -0.18 to 0.07 | -0.17 to 0.01 | -0.10 to 0.08 | -0.11 to 0.06 | — |

Table 2: This table compares how well each dimension reduction method performs in terms of logistic regression classifier AUC. Each entry shows the 95% confidence interval (CI) for the median of the differences between each method, computed using the Wilcoxon Rank-Sum test (where CI units measure the difference in AUC). An all-positive or all-negative confidence interval suggests that the difference is significant at an unadjusted p-value of $p < 0.05$.

| | top2pca vs. | top3pca vs. | top2pra vs. | top3pra vs. | top2pba vs. | top3pba vs. | top3amalg vs. | top4amalg vs. | top2slr vs. | top3slr vs. |
|---|---|---|---|---|---|---|---|---|---|---|
| top2pca | — | -0.02 to 0.03 | -0.0611 to -0.0003 | -0.0609 to -0.0005 | -0.0533 to -0.0004 | -0.045 to 0.005 | -0.005 to 0.046 | -0.004 to 0.049 | -0.002 to 0.051 | 0.005 to 0.059 |
| top3pca | -0.03 to 0.02 | — | -0.069 to -0.008 | -0.068 to -0.008 | -0.064 to -0.009 | -0.056 to -0.002 | -0.01 to 0.04 | -0.01 to 0.04 | -0.01 to 0.04 | -0.001 to 0.049 |
| top2pra | 0.0003 to 0.0611 | 0.008 to 0.069 | — | -0.03 to 0.03 | -0.03 to 0.04 | -0.02 to 0.05 | 0.02 to 0.08 | 0.02 to 0.08 | 0.03 to 0.08 | 0.03 to 0.09 |
| top3pra | 0.0005 to 0.0609 | 0.008 to 0.068 | -0.03 to 0.03 | — | -0.03 to 0.04 | -0.02 to 0.04 | 0.02 to 0.08 | 0.02 to 0.08 | 0.02 to 0.08 | 0.03 to 0.09 |
| top2pba | 0.0004 to 0.0533 | 0.009 to 0.064 | -0.04 to 0.03 | -0.04 to 0.03 | — | -0.02 to 0.03 | 0.02 to 0.08 | 0.02 to 0.08 | 0.02 to 0.08 | 0.03 to 0.09 |
| top3pba | -0.005 to 0.045 | 0.002 to 0.056 | -0.05 to 0.02 | -0.04 to 0.02 | -0.03 to 0.02 | — | 0.009 to 0.069 | 0.008 to 0.072 | 0.01 to 0.07 | 0.02 to 0.08 |
| top3amalg | -0.046 to 0.005 | -0.04 to 0.01 | -0.08 to -0.02 | -0.08 to -0.02 | -0.08 to -0.02 | -0.069 to -0.009 | — | -0.02 to 0.03 | -0.02 to 0.03 | -0.01 to 0.04 |
| top4amalg | -0.049 to 0.004 | -0.04 to 0.01 | -0.08 to -0.02 | -0.08 to -0.02 | -0.08 to -0.02 | -0.072 to -0.008 | -0.03 to 0.02 | — | -0.02 to 0.03 | -0.01 to 0.03 |
| top2slr | -0.051 to 0.002 | -0.04 to 0.01 | -0.08 to -0.03 | -0.08 to -0.02 | -0.08 to -0.02 | -0.07 to -0.01 | -0.03 to 0.02 | -0.03 to 0.02 | — | -0.01 to 0.03 |
| top3slr | -0.059 to -0.005 | -0.049 to 0.001 | -0.09 to -0.03 | -0.09 to -0.03 | -0.09 to -0.03 | -0.08 to -0.02 | -0.04 to 0.01 | -0.03 to 0.01 | -0.03 to 0.01 | — |
